# O-Glycosylated RNA Identification and Site-specific Prediction by Solid-phase Chemoenzymatic TnORNA method and PONglyRNA tool

**DOI:** 10.1101/2024.06.18.599663

**Authors:** Jiajia Li, Linshu Wang, Yan Chen, Shaomei Zhang, Zhongmin Wen, Xuechu Zhen, Haiyun Zhang, Yuan Zhou, Longjiang Xu, Shuang Yang

**Author notes:** Correspondence (S. Yang), (L. Xu), and (Y. Zhou). These authors equally contribute to this work (Jiajia Li, Linshu Wang).

## Abstract

Recent studies have shown that the cell surface undergoes post-transcriptional modification by N-linked glycosylation. However, the question of whether RNA can be glycosylated by O-glycans remains to be explored. The presence of O-glycosylation in cells is indirectly revealed by the presence of O-glycans on RNAs following treatment with O-glycoproteases. To identify RNA O-glycosylation, we have developed a chemoenzymatic method for capturing and enriching O-glycosylated RNA (O-glycoRNA) using covalent immobilization on a solid support. GalNAcEXO selectively releases Tn-containing O-glycosylated RNAs (TnORNA). Using this method and SPCgRNA, we compared the expression of O-glycoRNAs and N-glycoRNAs in pancreatic cancer cell lines and tissues. We found that glycosylated miR-103a-3p, miR-122-5p, and miR-4492 regulate pancreatic cancer cell growth and proliferation through the PI3K-Akt pathway. In vitro assays and PDAC tissue analysis confirmed the potential regulatory roles of Tn-O-glycosylated miRNAs in pancreatic tumor growth and metastasis. Furthermore, a significant number (131) of miRNAs carrying both N- and Tn-O-glycosylation were identified, indicating the co-occurrence of N-linked and O-linked glycosylation on small RNAs. We have also developed PONglyRNA, an online bioinformatic tool for the site-specific prediction of RNA glycosylation. PONglyRNA identifies glycosylation motifs based on RNA sequence and has been validated using our glycoRNA data. In conclusion, this study establishes robust experimental and computational tools for identifying O-linked glycoRNAs. Additionally, it uncovers the novel role of glycosylation in PDAC development and progression through altered glycosylation of oncogenic miRNAs.

**Graphical Abstract:** 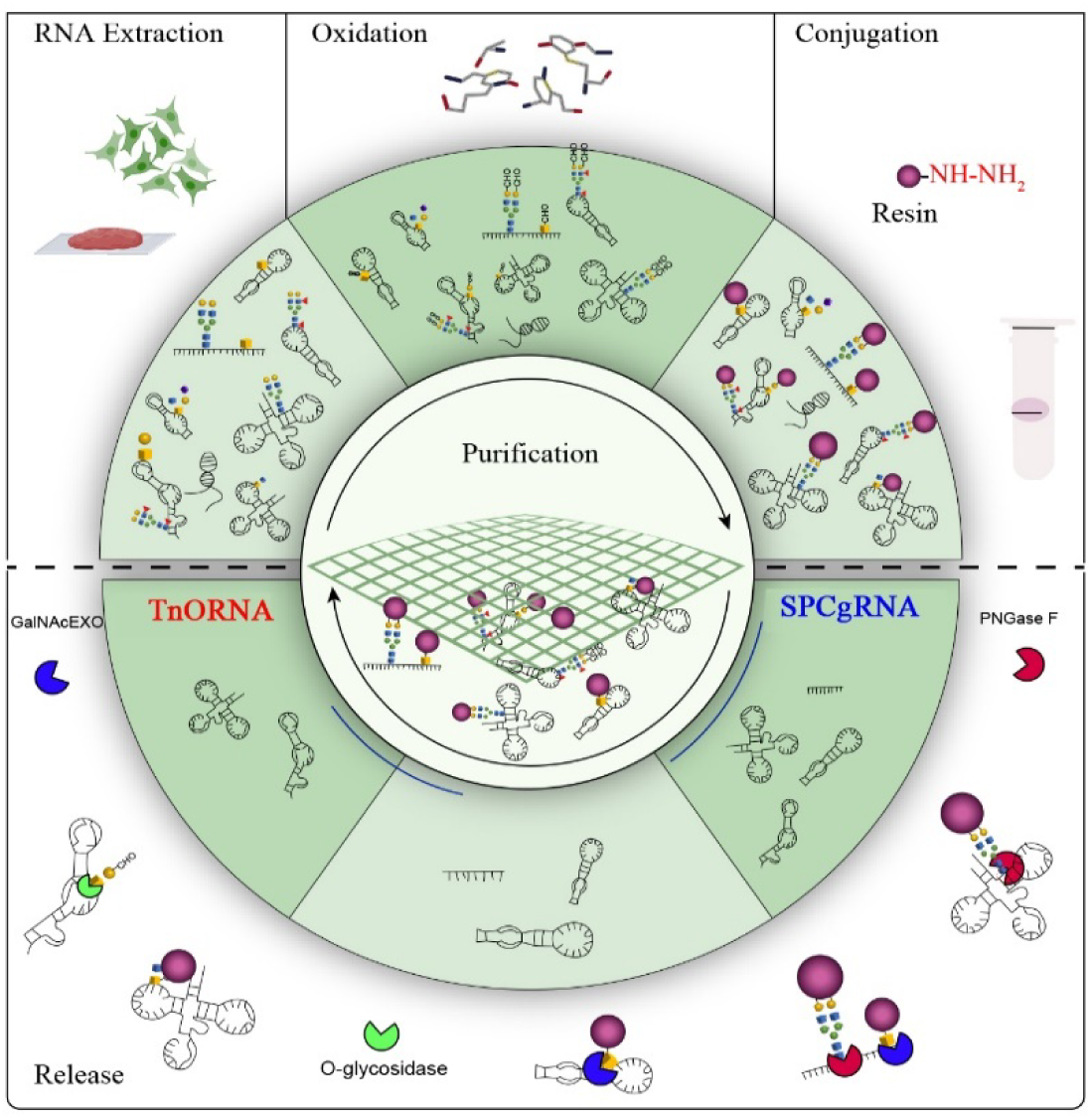

## Introduction

Diverse glycans and their linkages to other biological molecules constitute the complex glycome of a mammalian cell. Through glycosylation, glycans establish direct chemical bond linkages to various biological molecules, unleashing their unique and crucial regulatory roles in health and diseases, including but not limited to the wide involvements in inflammation ^1^, immune response evasion of viruses ^2^, and even regulation of kidney function ^3^. Glycomic profiling continues to identify new substrates of glycosylation, revealing intriguing and unique functions of glycosylation in both normal and disease states. On the other hand, glycosylation and the final glycan structures are determined not only by the activity of glycosyltransferases and glycosidases, but also by their subcellular localization and competitive interactions with other molecules, such as chaperones and protein complexes ^4^. Therefore, comprehensive profiling of the glycome is crucial to understand the multifactorial mechanisms of cellular glycan formation.

For several decades, glycosylation was thought to be exclusive to proteins and lipids. However, the last few years have witnessed a significant expansion in the known targets of glycosylation, with a prime example being the discovery of glycosylated RNA (glycoRNA). Recent studies have revealed that mammalian cell surface RNAs, particularly small nucleolar RNAs (snoRNAs), can undergo N-linked glycosylation (N-glycosylation) ^5^. These nuclear-genome encoded RNAs (ngRNAs) reach the cell surface through various mechanisms. One pathway involves forming ribonucleoprotein (RNP) complexes with transmembrane proteins after transcription. Additionally, studies suggest that certain ngRNAs can directly bind to membrane lipids in humans, offering another route for surface association ^6^. The presence of sialic acid structures on these glycoRNAs opens new avenue for cell communication and immune regulation. These glycans might interact with immune system players like Siglecs or galectins, potentially influencing the precision of the immune response ^5^. Scientists have further identified a role for cell surface RNAs, transported by mammalian homologs of sid-1, in neutrophil recruitment and adhesion ^7^. These P-selectin-binding glycoRNAs specifically influence interactions between neutrophils and endothelial cells, highlighting a novel role for RNA in cellular communication.

Despite significant progress, our understanding of RNA glycosylation lags behind that of protein glycosylation. O-glycosylation, a major category in protein modification, remains poorly understood in the context of RNA. Deregulated protein O-glycosylation, particularly the presence of truncated forms (T or Tn antigens), is a hallmark of various diseases, including breast cancer^8^, lung cancer ^9^, and pancreatic ductal adenocarcinoma (PDAC) ^10^. In healthy pancreases, glycoproteins play essential roles like protecting ducts, while PDAC exhibits dysregulated glycosylation with aberrant glycans linked to disease progression. The PDAC glycome shows increased sialylation, fucosylation, and truncated O-glycans (Tn and sTn antigens) ^10–12^. However, the existence and precise nature of O-glycosylation in RNAs, especially in PDAC, remain open questions demanding further investigation.

Contrary to the widespread glycosylation observed across diverse protein families, only a limited number of RNA categories have been identified to undergo glycosylation. However, non-coding RNAs (ncRNAs), particularly microRNAs (miRNAs), are functionally associated with this process. These miRNAs can regulate N- and O-glycan biosynthesis at multiple stages, influencing the overall glycome ^13,14^. Kurcon et al. pioneered the use of miR-200f, a known epithelial-to-mesenchymal transition (EMT) regulator, to identify EMT-associated glycoenzymes such as *ST3GAL5* and *ST6GALNAC5*. Their study highlights the potential of miRNAs as tools to understand how ncRNAs control glycosylation ^15,16^. These ncRNA-regulated glycoenzymes are involved in various steps of glycan biosynthesis, including initiation, core-extension, elongation, and capping. For example, miR-98-5p potentially downregulates *ALG3* (α1,3-mannosyltransferase), contributing to non-small cell lung cancer (NSCLC) malignancy ^17^. Similarly, miR-199a dysregulates *ST6GAL1* expression in cancer, ultimately impacting the *ERBB2*/*ERBB3* signaling pathway ^18^. Bisecting-GlcNAc, a hallmark of cancer, is produced by the enzyme *MGAT3*. However, *MGAT3* is often downregulated in tumors, leading to reduced cell-cell adhesion. Conversely, miR-23a activation promotes hepatoma cell metastasis by targeting *MGAT3* ^19^. Interestingly, miR-23b targets *MGAT3* on the Tau protein, potentially offering a therapeutic approach for Alzheimer’s disease (AD) by influencing the Akt/GSK-3b pathway ^20^.

The relationship between miRNAs and Polypeptide N-acetylgalactosaminyltransferases (GALNTs) is particularly intricate, demonstrating context-dependent effects on cancer progression ^21^. This complex interplay underscores the importance of understanding which specific GALNT substrates are targeted by miRNAs in different contexts. miRNA networks offer a promising avenue to unravel these regulatory pathways. For example, miR-214 upregulates *GALNT7*, leading to suppressed aggressiveness in cervical cancer cells ^22^. Conversely, miR-17 downregulates *GALNT7*, promoting tumor growth in hepatocellular carcinoma ^23^. Melanoma presents another layer of complexity, as miR-30b/30d downregulates *GALNT7*, weakening the immune response ^24^. These seemingly contradictory findings across different cancers highlight the critical need for context-specific analysis.

PDAC is also characterized by the dysregulation of numerous miRNAs, significantly impacting various aspects of cancer progression. Certain miRNAs, like miR-221/222 and miR-210, promote invasion and cell survival, while others, such as miR-100 and Let-7, function as tumor suppressors by inhibiting cell proliferation and invasion. This intricate interplay between miRNAs and their targets underscores their potential as therapeutic targets in the fight against PDAC ^25^. However, crucial questions remain: 1) Do miRNAs undergo O-glycosylation, and can altered glycoenzyme expression impact this process and its consequences? 2) While enriched N-glycosylation has been identified in snoRNAs, it is intriguing to speculate whether miRNAs, as a distinct ncRNA category, might exhibit a preference for O-glycosylation. 3) Is truncated O-glycosylation differentially expressed in PDAC?

The lack of efficient methods for detecting glycoRNA has become the main bottleneck to systematically profiling the RNA glycome. In response, detecting RNA glycosylation has emerged as an active research area with several promising methods under development. While evidence supports N-glycan modifications on RNAs, techniques targeting the glycans themselves provide the most reliable confirmation. One approach focuses on sialic acids, essential components of N-glycans. These require activation by specific enzymes (CMP-Sia synthases) before incorporation onto RNA molecules. Pioneering work by Bertozzi and colleagues employed metabolic labeling with azide-tagged sugars, commonly used for protein and lipid glycan analysis. This work revealed that these tags can also label RNA molecules, suggesting the presence of RNA modifications with sugars ^5^. This technique was further refined for high-resolution imaging of single-cell glycoRNA. It employs a combined approach: aptamer-based enrichment and RNA in situ hybridization-mediated proximity ligation assay (ARPLA) ^26^. This method’s sensitivity and specificity allowed the researchers to reveal the location, dynamics, and potential roles of glycoRNA in cancer and immune response, showcasing its value for studying these unique molecules and their functions. Another approach utilizes the enzyme galactose oxidase (GAO) to specifically target N-glycosylated RNAs. The resulting oxidized structures can then be conjugated to hydrazide beads, capturing N-glycoRNAs on a solid phase while leaving non-glycosylated RNAs in the supernatant ^27^. While techniques like PNGase F treatment and metabolic labeling-click chemistry have identified glycosylated miRNAs and other small RNAs in pancreatic cancer, they lack the necessary specificity for O-glycans ^5,27^. Even sialic acid metabolic labeling, although it might capture some O-glycosylated transcripts, suffers from limitations in specific detection.

To address this challenge, we introduce a robust solid-phase chemoenzymatic method for specifically enriching glycosylated RNAs, followed by isolation of O-glycoRNAs. This method builds upon previous work demonstrating the selective oxidation of Gal/GalNAc residues on RNAs by galactose oxidase (GAO) and subsequent conjugation to hydrazide resin ^27^. This two-step approach ensures the capture of only glycosylated RNAs during TRIzol extraction by eliminating glycoproteins beforehand. This method enriches for O-glycoRNAs through a chemical conjugation step. Crucially, specific O-glycosidases, such as O-glycosidase or GalNAcEXO, can then cleave O-glycoRNAs from the resin. Notably, GalNAcEXO allows for the selective enrichment of **<u>Tn</u>**-containing **<u>O</u>**-glyco**<u>RNA</u>**s (TnORNA). Using this approach, we analyzed the profile of O-glycosylated RNAs containing Tn (Tn-O-glycoRNAs), particularly glycosylated miRNAs, in pancreatic ductal adenocarcinoma (PDAC) cells and tissues. Building on the identified glycoRNAs, we further developed PONglyRNA (**<u>P</u>**redictor of **<u>O</u>**- and **<u>N</u>**-linked **<u>gly</u>**co**<u>RNA</u>**), a bioinformatic tool that predicts both N-linked and O-linked RNA glycosylation at specific sites. This study not only establishes robust experimental and computational tools for comprehensively identifying glycoRNAs but also aims to decipher the potential functional roles of RNA glycosylation in PDAC development and progression.

## Results

### GalNAcEXO possesses the potential to specifically target and distinguish O-glycoRNAs containing the Tn antigen

**Table 1** summarizes several existing techniques that target glycoRNAs. Metabolic labeling combined with click chemistry or ARPLA relies on the incorporation of CMP-sialic acid during cellular metabolism. However, ARPLA cannot differentiate between N-linked and O-linked sialic acid-containing glycoRNAs ^5,26^. While metabolic labeling methods coupled with O-glycoproteases (StcE or OpeRATOR) offer potential for identifying RNA O-glycosylation, their ability to specifically cleave RNA nucleotide bonds remains unclear. These O-glycoproteases primarily function as endopeptidases, targeting amino acid bonds in proteins ^28–30^. The SPCgRNA method involves immobilizing Gal- or GalNAc-containing glycoRNAs, followed by their release with PNGase F for N-glycosylation analysis ^27^. We hypothesize that, building upon this approach, O-glycosidases can cleave the glycosidic bond between Gal or GalNAc and the RNA backbone. This is based on the assumed functional similarity between O-glycosidases acting on glycoproteins and RNA. Consequently, these enzymes could be used to cleave O-glycoRNAs harboring the T or Tn antigen.

**Table 1.**
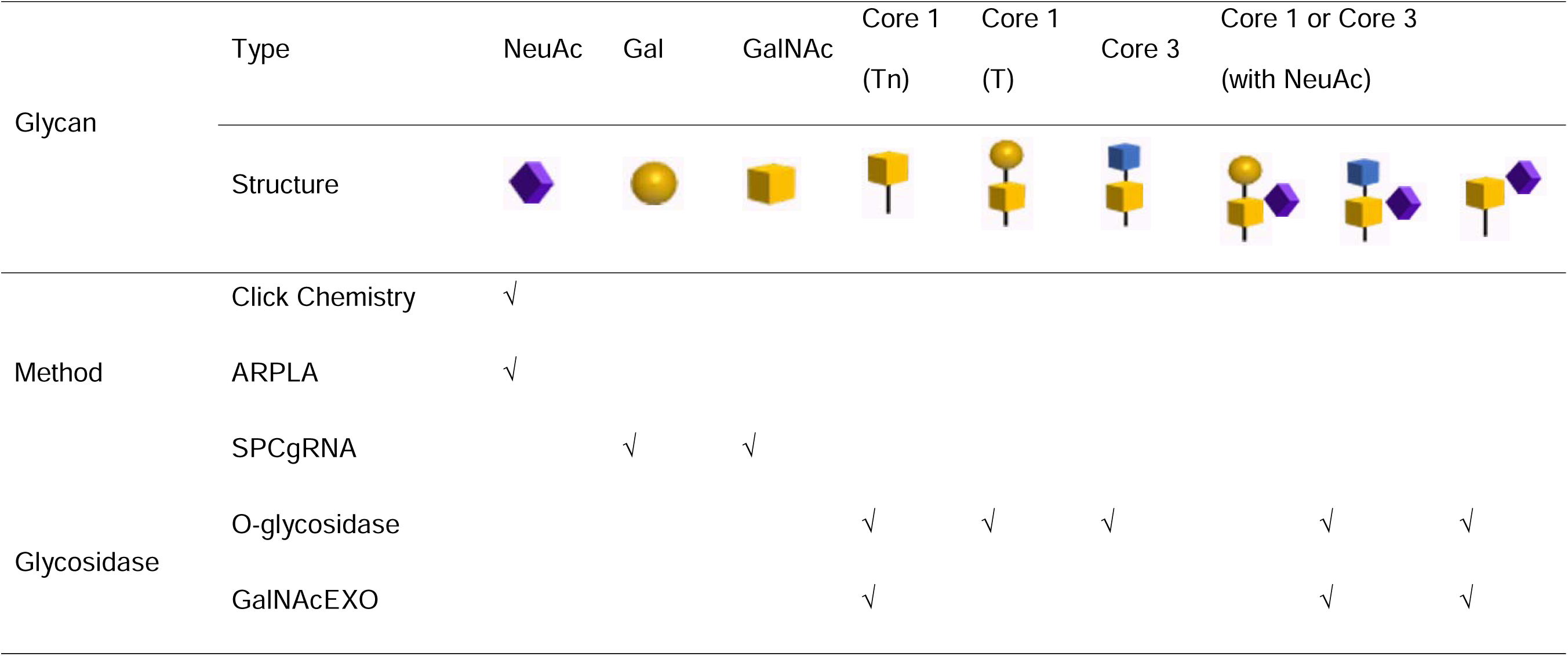
Comparison of methods and glycosidases for glycoRNA identification. This table compares different methods and glycosidases used for identifying glycoRNAs. PNGase F effectively hydrolyzes N-glycoRNAs using the SPCgRNA method, while its counterpart for O-glycoRNAs, such as O-glycosidase or GalNAcEXO, can be employed for O-glycoRNA enrichment and subsequent identification. Footnote: √ = active; NeuAc = sialic acid; Gal = Galactose; GalNAc = N-Acetylgalactosamine; Tn and T are two antigens.

**Table 2.**
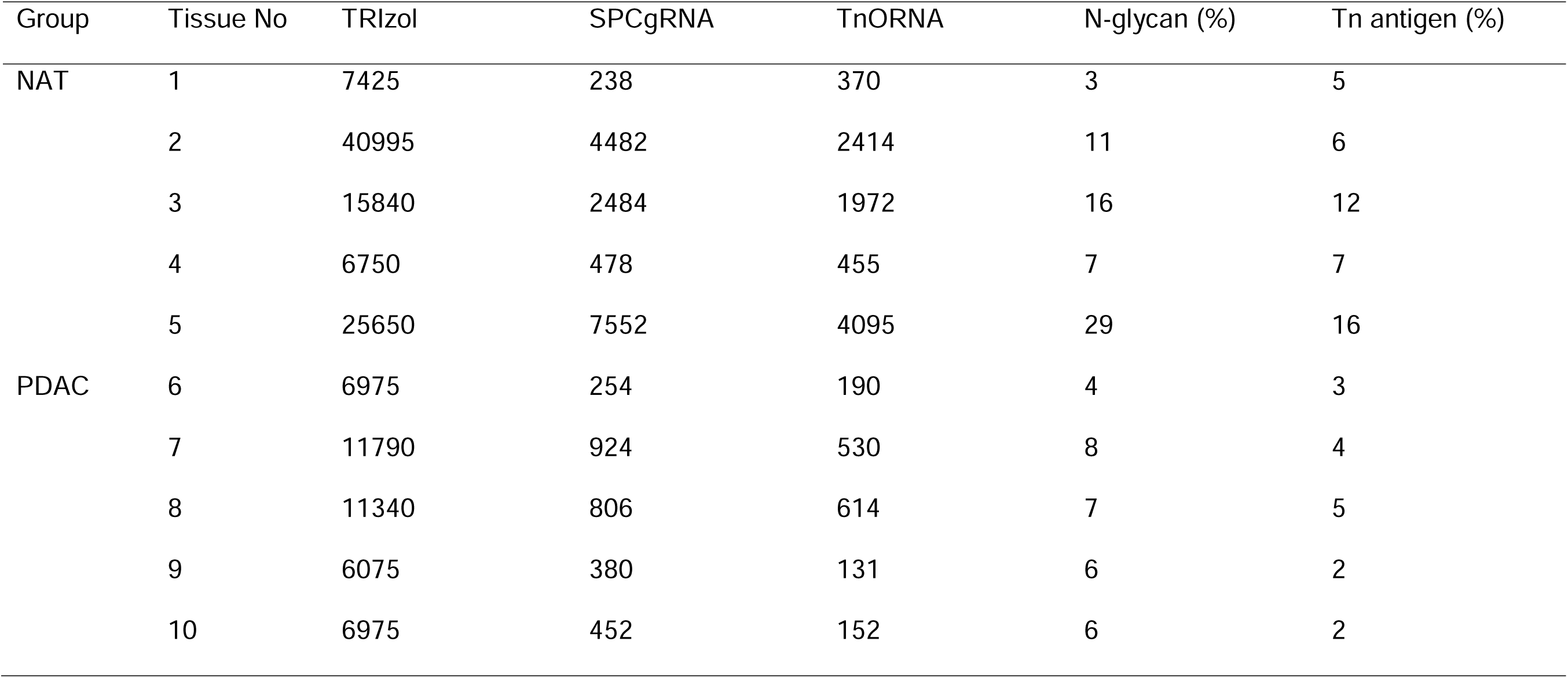
Comparison of glycoRNA content in PDAC and NAT tissue clinical samples using SPCgRNA and TnORNA methods. This table presents the results of quantifying glycoRNA content in paired pancreatic ductal adenocarcinoma (PDAC) and normal adjacent tissue (NAT) tissue samples from clinical datasets. The glycoRNA content was determined using two methods: the standard SPCgRNA method and the recently developed TnORNA method, which specifically enriches for Tn-motif containing O-glycoRNAs.

| Group | Tissue No | TRIzol | SPCgRNA | TnORNA | N-glycan (%) | Tn antigen (%) |
| --- | --- | --- | --- | --- | --- | --- |
| NAT | 1 | 7425 | 238 | 370 | 3 | 5 |
|  | 2 | 40995 | 4482 | 2414 | 11 | 6 |
|  | 3 | 15840 | 2484 | 1972 | 16 | 12 |
|  | 4 | 6750 | 478 | 455 | 7 | 7 |
|  | 5 | 25650 | 7552 | 4095 | 29 | 16 |
| PDAC | 6 | 6975 | 254 | 190 | 4 | 3 |
|  | 7 | 11790 | 924 | 530 | 8 | 4 |
|  | 8 | 11340 | 806 | 614 | 7 | 5 |
|  | 9 | 6075 | 380 | 131 | 6 | 2 |
|  | 10 | 6975 | 452 | 152 | 6 | 2 |

### TnORNA effectively enriches Tn-containing O-glycoRNAs through GalNAc residue digestion

The TnORNA method specifically enriches O-glycoRNAs harboring terminal GalNAc (Tn) modifications by releasing RNAs from the solid-phase using GalNAcEXO, as detailed in **Figure 1A**. The workflow includes: (**1**) Total RNA Extraction: First, total RNA is extracted from cells or tissues using TRIzol to remove proteins. Protein glycosylation can interfere with identifying RNA glycosylation ^31^. (**2**) RNA Oxidation and Immobilization: The extracted RNA is then oxidized with GAO and immobilized on a hydrazide resin. (**3**) Washing and Release: After washing with DEPC water to remove unbound molecules, the immobilized RNA is released by treatment with either GalNAcEXO (which specifically cleaves GalNAc residues) or another O-glycosidase capable of cleaving the Gal-GalNAc disaccharide. (**4**) Validation: The effectiveness of TnORNA was validated using controlled experiments with a pancreas cell line (MIA PaCa-2) (**Figure 1B**). Four fractions were collected for RNA identification and quantification: #1 (total RNA), #2 (wash), #3 (supernatant after GalNAcEXO digestion), and #4 (supernatant after O-glycosidase digestion). The absence of RNA in the washing solution (fraction 2) confirms the complete removal of unbound RNA. Conversely, the presence of RNA in fractions 3 and 4 (around 10% of the total) suggests successful cleavage of targeted O-glycoRNAs by the respective enzymes (GalNAcEXO for fraction 3 and potentially another O-glycosidase for fraction 4). This cleavage likely occurs through hydrolysis of the glycosidic bond between the sugar and the RNA backbone.

**Figure 1.**
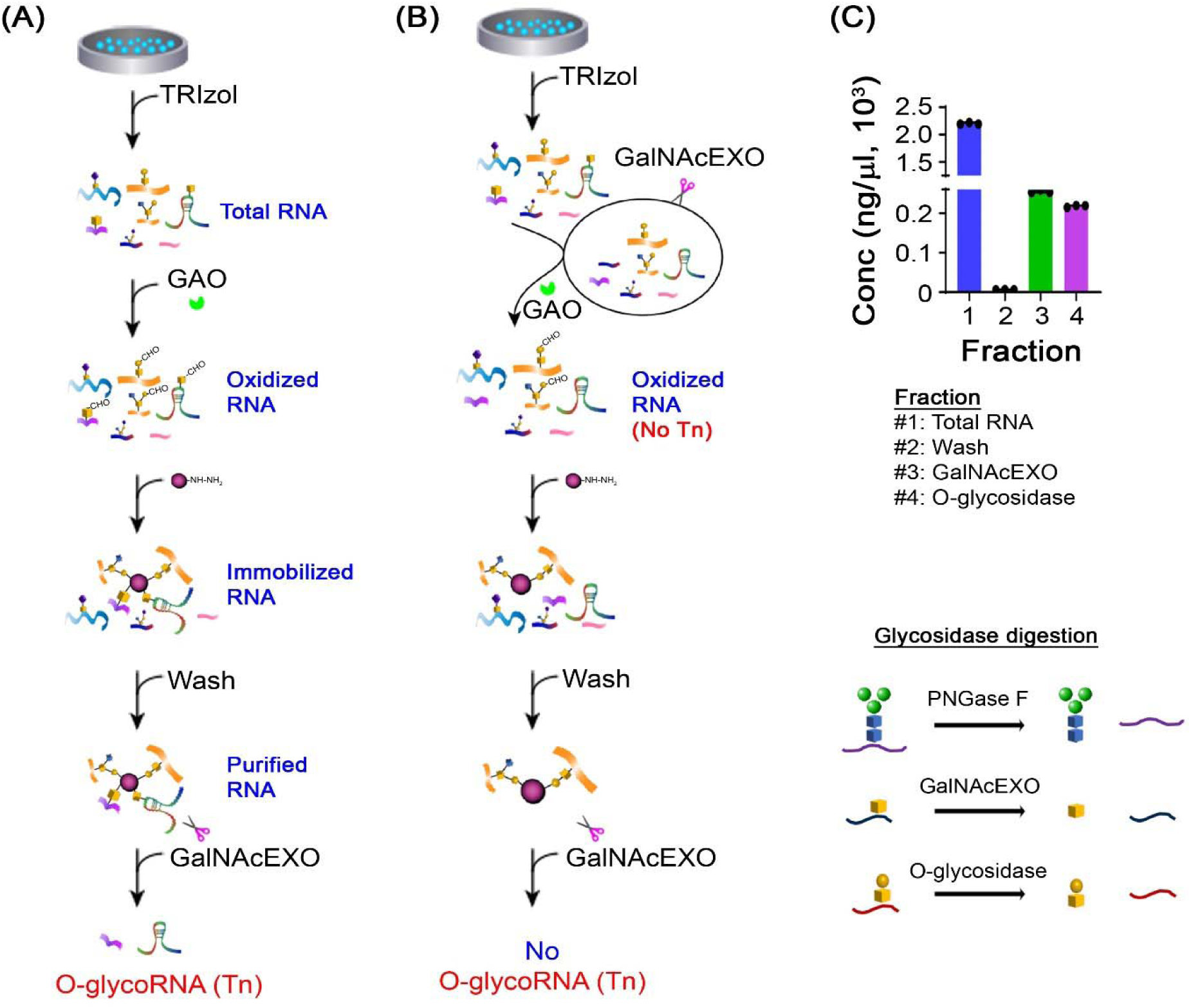
TnORNA: a method for specific enrichment of Tn-containing O-glycoRNAs. (A) TnORNA workflow schematic. This schematic illustrates the key steps of the TnORNA method: total RNA extraction, galactose oxidation of glycoRNAs, covalent conjugation, washing and elution, GalNAcEXO digestion. (B) O-glycoRNA enrichment comparison using sequential GalNAcEXO and O-glycosidase. The X-axis represents different fractions including total RNA (starting material containing all RNA species), washed RNA after covalent conjugation (RNA remaining after unbound molecules are removed following conjugation to hydrazide resin beads), Tn-containing O-glycoRNA released by GalNAcEXO, T-containing O-glycoRNA released by additional O-glycosidase digestion. (C) Effect of GalNAcEXO pretreatment. When GalNAcEXO digestion is performed before GAO oxidation (pre-GalNAcEXO), it cleaves Tn-containing O-glycans beforehand. Consequently, no RNA is expected to be released by the subsequent GalNAcEXO step.

### Optimizing TnORNA parameters for improved O-glycoRNA identification

To confirm the presence of Tn-containing O-glycoRNAs in fraction #3, we pretreated the total RNA with GalNAcEXO before GAO oxidation **(Figure 1C**). This pretreatment was designed to remove all terminal GalNAc residues (Tn modifications) from O-glycoRNAs. Consequently, we expected no detectable glycosylated RNA in fraction #3 after GalNAcEXO pretreatment. However, a significant amount of O-glycoRNA remained in the MIA PaCa-2 cell line following this pretreatment, which contradicts our prediction based on **Figure 1C**. To elucidate the unexpected persistence of O-glycoRNAs after GalNAcEXO pretreatment, we explored two potential mechanisms. One hypothesis was that sialic acids or glycoproteins might compete with RNA for binding to the galactose oxidase (GAO) enzyme, hindering the oxidation of GalNAc residues on the RNA backbone. Alternatively, an excessive amount of RNA could overwhelm GalNAcEXO’s capacity, leaving some Tn modifications undigested. To investigate these possibilities, we first pretreated total RNA with SiaEXO to remove sialic acids (**Figure 2A**). We then examined the effect of adding MUC5AC, a known Tn-containing glycoprotein, to the RNA before GalNAcEXO treatment (**Figure 2B**). Our results demonstrate a decrease in detectable Tn-containing O-glycoRNAs after sialic acid removal. Conversely, there is a significant increase in detectable Tn-containing O-glycoRNAs in cases with more MUC5AC interference. These findings suggest that an excessive amount of RNA can indeed hinder GalNAcEXO activity.

**Figure 2.**
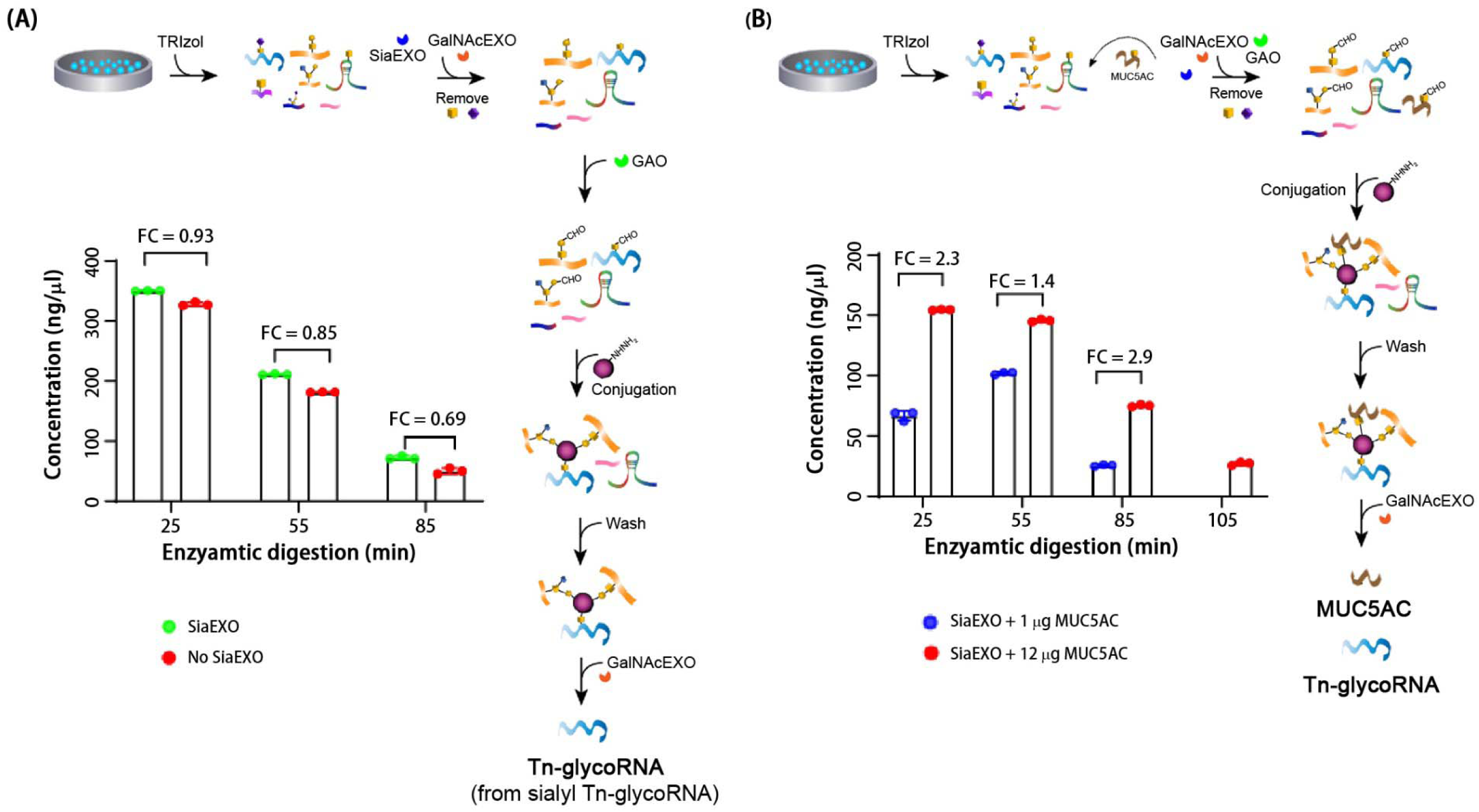
Effects of sialic acids and mucin-type O-glycopeptides on Tn-containing O-glycoRNA digestion after GalNAcEXO pretreatment. (A) Sialic acid effect: incomplete Tn-O-glycoRNA digestion might be due to GAO consumption by sialic acid oxidation, converting sTn to Tn. Sialidase (SiaEXO) is used to remove sialic acids prior to GAO oxidation. The amount of released RNA has no significant difference before and after sialic acid removal, suggesting that sialic acids are not responsible for incomplete O-glycoRNA release. (B) Mucin-type O-glycopeptides to explore the interference of proteins or degraded peptides mixed with total RNA: as elution time increases, the RNA concentration obtained with higher MUC5AC (mucin) content significantly differs from that obtained with lower MUC5AC. This indicates that proteins within the sample affect GalNAcEXO enzyme digestion efficiency and are the primary cause of incomplete O-glycoRNA digestion.

Extending GalNAcEXO digestion time to 85 minutes yielded a further increase in O-glycoRNAs, indicating that longer enzyme treatment can process more Tn-containing RNA substrates (**Figure S1**). Conversely, increasing the concentration of MUC5AC, a Tn-containing glycoprotein competitor, from 1 to 12 μg with the same GalNAcEXO treatment and duration resulted in a lower final concentration of O-glycoRNAs (**Figure S1**). This suggests that higher concentrations of Tn-containing glycoproteins compete with target RNAs for GalNAcEXO activity. Taken together, these results demonstrate that both sialic acids and abundant Tn-containing glycoproteins can impede the efficiency of GalNAcEXO pretreatment during TnORNA analysis. Therefore, minimizing glycoprotein contamination during RNA extraction using TRIzol is crucial for accurate O-glycoRNA analysis.

We investigated the optimal GalNAcEXO amount for complete Tn-containing O-glycoRNA digestion. We treated varying starting RNA amounts (20, 40, and 60 μg) with a fixed GalNAcEXO concentration (20 μg) (**Figure S1**). While no significant difference in O-glycoRNA concentration was observed between control and GalNAcEXO-treated samples at the highest RNA amount (60 μg), a substantial amount of O-glycoRNAs remained at 40 μg. Notably, using the TnORNA method with Nanodrop measurement, pretreatment with 20 μg GalNAcEXO effectively digested all detectable O-glycoRNAs in the lowest RNA amount (20 μg). These findings suggest that a 1:1 ratio of GalNAcEXO to RNA might be ideal for complete digestion.

### TnORNA analysis of glycoRNAs from PDAC cell lines

The TnORNA method was employed to enrich Tn-containing O-glycoRNAs from A549 and MIA PaCa-2 PDAC cell lines. The amount of GalNAcEXO used was proportionally adjusted to the total RNA quantity (**Figures 3A** & **S1**). This ensures complete removal of the Tn antigen during pretreatment, which is critical to prevent underestimating the number of Tn-containing O-glycoRNAs, especially for less abundant species. While optimizing GalNAcEXO usage is essential for maximizing O-glycoRNA identification, it does not diminish the effectiveness of TnORNA. Despite potential limitations, this method successfully identified hundreds of Tn-O-glycoRNAs. Notably, tRNAs and rRNAs were the most abundant O-glycoRNA species detected (**Figures 3B** and **3C**). Correlation analysis using the Pearson correlation coefficient revealed strong and significant correlations between the O-glycoRNA expression profiles of control and experimental groups in the cell lines (**Figure 3D**). This suggests that the TnORNA method’s pretreatment, likely due to the optimized GalNAcEXO usage, effectively minimizes background noise from O-glycoproteins and non-Tn-containing O-glycoRNAs.

**Figure 3.**
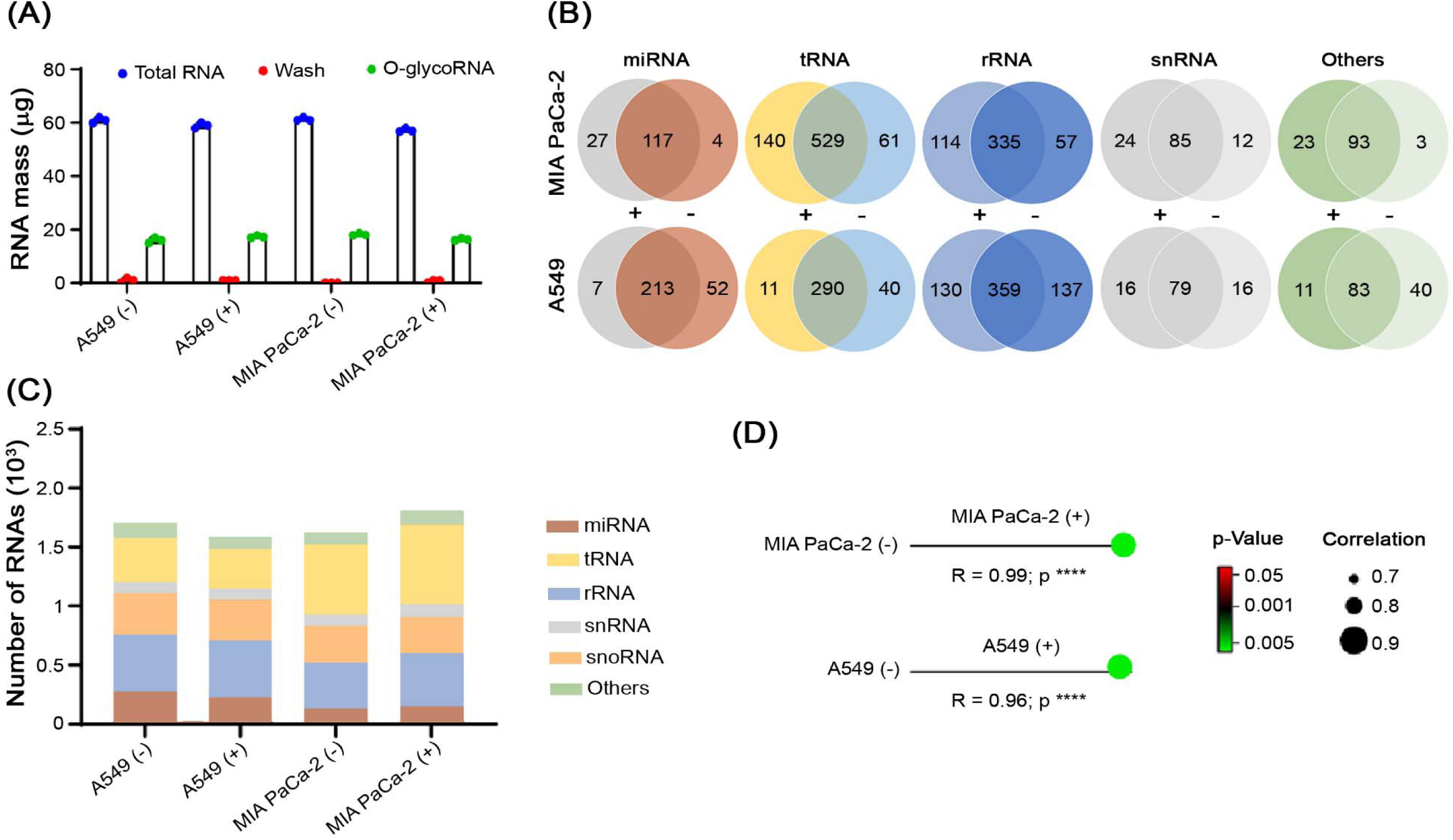
Identification of O-glycoRNAs from A549 and MIA PaCa-2 cell lines using the TnORNA method. (A) Changes in RNA concentration are used in TnORNA for both A549 and MIA PaCa-2 cell lines. The data points include starting amount of RNA before treatment (initial total RNA), concentration of unbound RNA removed during the washing step (wash step cleaning solution), concentration of the collected eluent, enriched in O-glycoRNAs (final eluent). Samples are labeled as A549(-) (A549 cells treated with the standard TnORNA method), A549(+) (A549 cells treated with GalNAcEXO pretreatment), MIA PaCa-2(-) (MIA PaCa-2 cells treated with the standard TnORNA method), and MIA PaCa-2(+) (MIA PaCa-2 cells treated with GalNAcEXO pretreatment). (B) Tn-containing-O-glycoRNA identification. The specific types and quantities of Tn-O-glycoRNAs are identified by TnORNA. (C) Tn-O-glycoRNA comparison. The relative abundance of different Tn-O-glycoRNAs between the A549(-) and A549(+) groups, between the MIA PaCa-2(-) and MIA PaCa-2(+) groups are compared. (D) The correlation analysis shows the expression profiles of O-glycoRNAs between the pre-control and experimental groups. The Pearson correlation coefficient (R) is used to evaluate the strength and significance of the correlation. A strong, significant correlation between A549 and MIA PaCa-2 cells suggests that the pretreatment in the TnORNA method effectively reduces interference from GalNAcEXO or Tn-O-glycoRNA, possibly by increasing the amount of GalNAcEXO.

### PDAC tissues exhibit significantly higher abundance and distinct expression patterns of N-glycoRNAs compared to Tn-containing O-glycoRNAs

Previous studies have identified N-glycoRNAs in PDAC cell lines ^27^. However, O-glycoRNAs remained unexplored in this context. Here, we present the first analysis of Tn-containing O-glycoRNA profiles alongside N-glycoRNAs in PDAC tissues. Our data revealed that N-glycoRNAs were generally more abundant than Tn-containing O-glycoRNAs in both normal adjacent tissues (NATs) and PDAC tissues (**Figure 4A**). This difference likely stems from the specific enrichment of TnORNA, which targets only Tn-containing O-glycoRNAs since GalNAcEXO has no activity on other complex O-glycans (**Figure S2**). Our analysis revealed a significant increase in both N-glycoRNAs and Tn-containing O-glycoRNAs in PDAC tissues compared to NATs. PDAC tissues exhibited over fivefold enrichment of N-glycoRNAs (3046 vs. 563) and nearly sixfold enrichment of O-glycoRNAs (1861 vs. 323) compared to NATs. Interestingly, unlike PDAC cell lines, snoRNAs were the most abundant glycosylated RNA species in both tissues, followed by rRNAs and tRNAs (**Figure 4B**). Principal component analysis (PCA) provided further insights. Notably, certain RNAs clustered together in both NATs and PDACs for both N- and O-glycoRNAs (**Figure 4C**). This suggests the co-occurrence of N- and O-linked glycosylation on these specific small RNAs. Additionally, PCA confirmed distinct expression patterns of N-glycoRNAs and Tn-containing O-glycoRNAs between NATs and PDACs (**Figure 4D**).

**Figure 4.**
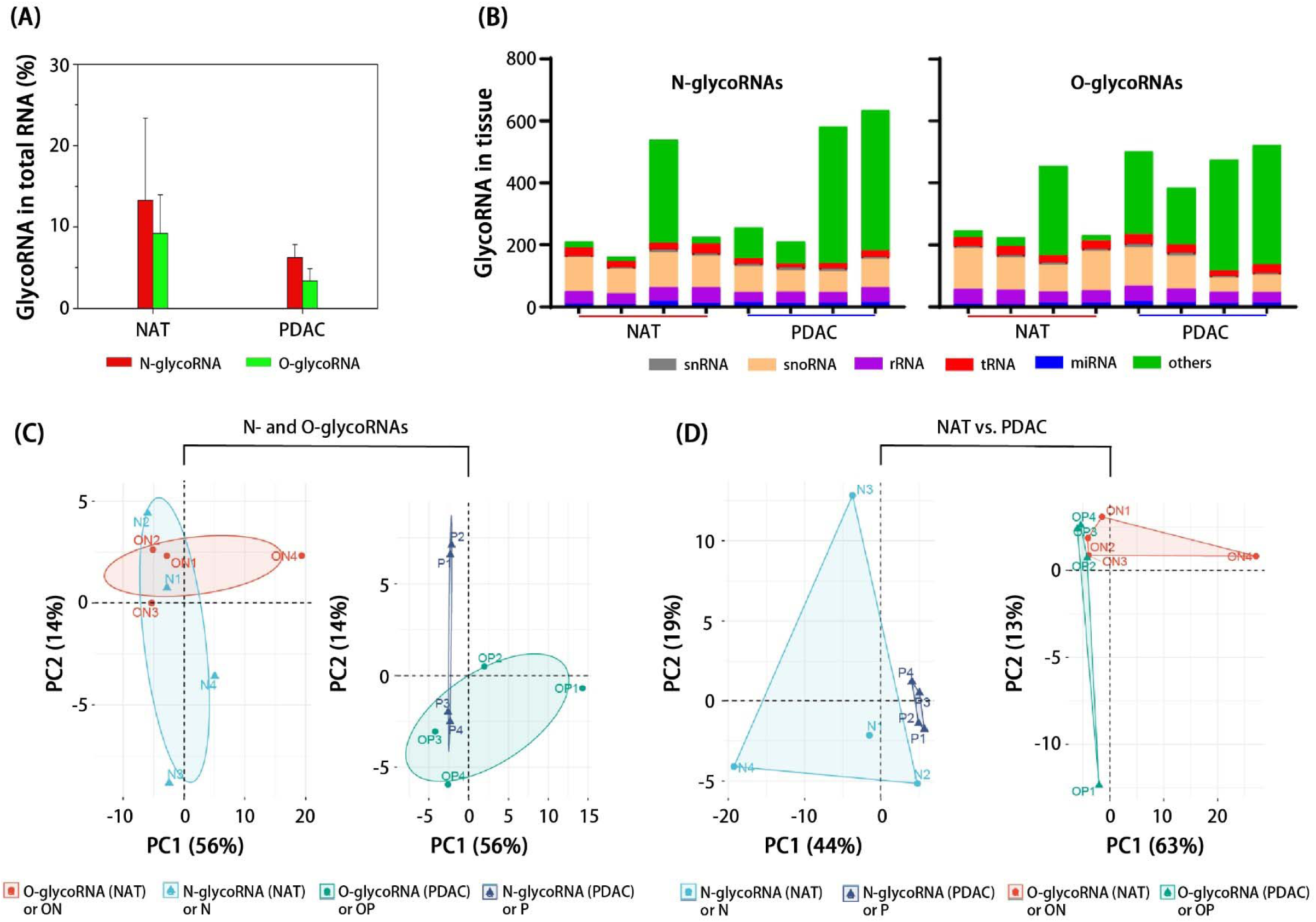
GlycoRNA expression in NAT and PDAC using SPCgRNA and TnORNA methods. (A) N- and O-glycoRNA identification in normal adjacent tissue (NAT) and pancreatic ductal adenocarcinoma (PDAC) tissue samples. (B) The different types of small glycoRNAs are identified in samples from 4 NAT and 4 PDAC patients. The number of glycoRNAs identified remains consistent across different individuals, unlike other types of small RNAs. (C) The principal component analysis (PCA) of N-glycoRNAs and O-glycoRNAs from NAT and PDAC samples. The PCA plot suggests that some small RNAs carry both N- and O-linked glycosylation, as indicated by their positions on the plot. (D) PCA analysis of N-glycoRNAs and O-glycoRNAs reveals the differential expression of glycoRNAs between NAT and PDAC tissues.

### Unique glycoRNA patterns in PDAC tissue implies altered miRNA regulation through glycosylation

Our analysis of miRNA profiles revealed distinct glycosylation patterns between PDAC tissues and NATs (**Figure 5A**). Notably, NATs displayed a higher abundance of O-glycosylated snoRNAs (90) compared to PDACs (60). Conversely, PDAC tissues showed a slight increase in Tn-containing O-glycosylated miRNAs (12) compared to NATs (8). Interestingly, both tissues had similar numbers of O-glycosylated rRNAs (>30). Furthermore, analysis of MIA PaCa-2 cells identified a significantly higher number of miRNAs carrying both N- and O-glycosylation (131) compared to those modified by only one type (40 with both). This suggests that PDAC tissues, with their greater complexity of glycans, might harbor a wider variety of miRNAs harboring both N- and O-glycosylation compared to cell lines.

**Figure 5.**
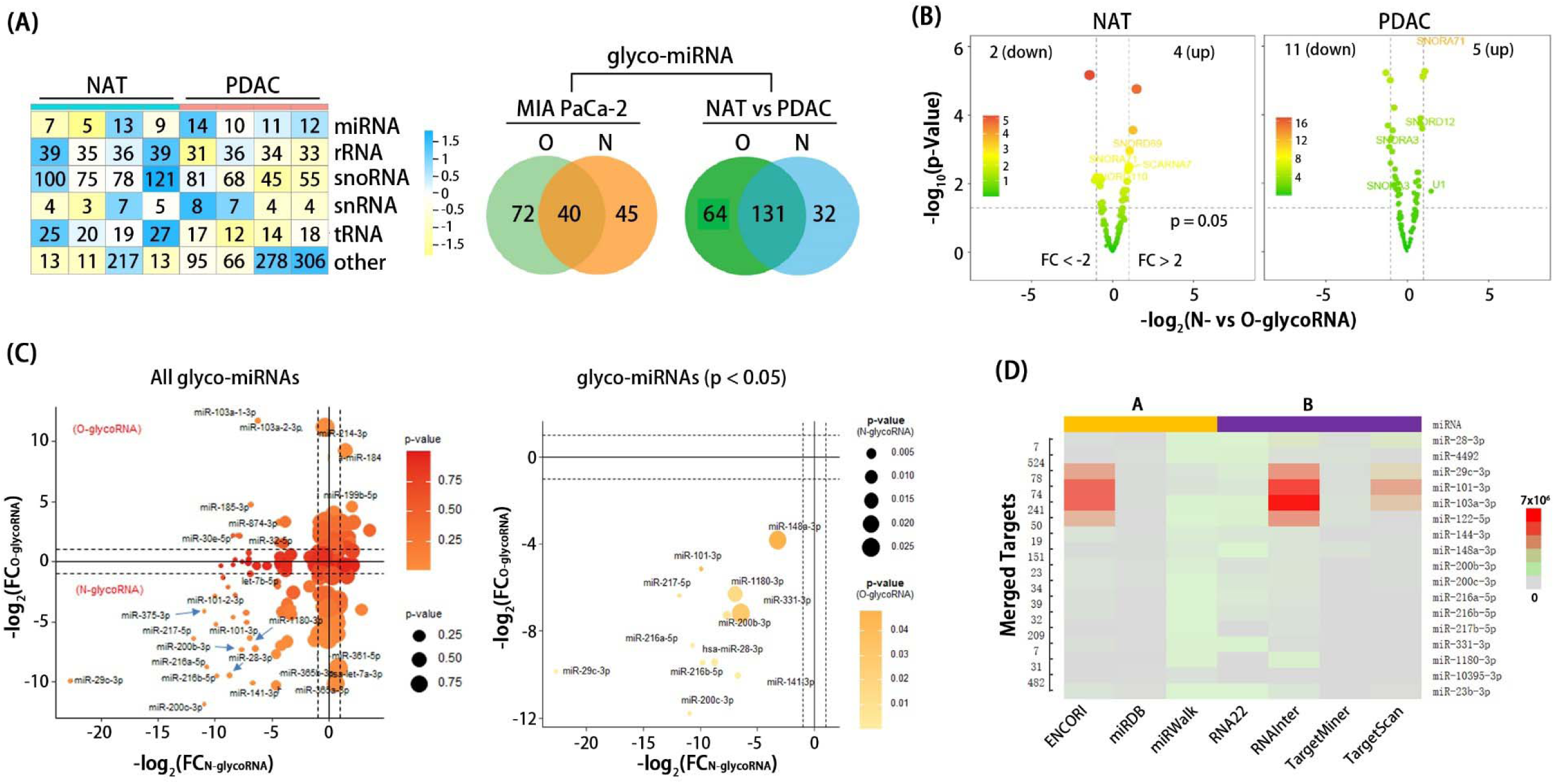
Glycosylated miRNAs identified from pancreatic cancer cell lines and tissues. (A) Shared N-glycosylated and O-glycosylated miRNAs from normal (NAT) and pancreatic cancer (PDAC) tissues. MIA PaCa-2 shows 40 shared miRNAs concurrently present in SPCgRNA and TnORNA. PDAC and NAT tissues identify 131 shared miRNAs. (B) The volcano plots show differential expression of the 131 shared N-glycoRNAs and O-glycoRNAs identified by comparing PDAC and NAT tissues. (C) Correlation of N- and O-glycosylation on shared miRNAs in scatter plots and correlation diagrams for miRNAs in 4 pairs of patients PDAC and NAT tissues. “FC_N-glycoRNA_” indicates the fold change of miRNAs by SPCgRNA, while “FC_O-glycoRNA_” represents the fold change by TnORNA. The correlation coefficient signifies that the same miRNA can be glycosylated by N-glycans and Tn antigen. MiRNA scatter plots show statistically significant (p < 0.05) and consistent expression in shared miRNAs between PDAC and NAT tissues from 4 pairs of patients. (D) Predicted downstream targets of shared and unique miRNAs from 7 miRNA target prediction websites (e.g., ENCORI, miRDB) to predict downstream target genes associated with the 13 significantly shared and 3 significantly unique miRNAs (**Table S1**).

Volcano plot analysis of glycosylated RNAs revealed distinct expression patterns between NATs and PDACs (**Figure 5B**). NATs displayed only a few differentially expressed glycosylated RNAs (4 upregulated and 2 downregulated), while PDACs harbored a larger number (5 upregulated and 11 downregulated), suggesting that these differentially expressed molecules in PDACs might play a role in tumor development. Furthermore, correlation analysis of miRNAs shared across four patient PDAC tissues provided another interesting finding. We analyzed these tissues using SPCgRNA and TnORNA. Results showed co-occurrence of N- and O-glycosylation on the same miRNAs (**Figure 5C**). Our analysis revealed a significant positive correlation between FC_N-glycoRNA_ (N-glycosylation enrichment) and FC_O-glycoRNA_ (Tn-glycosylation enrichment), indicating that these modifications can coexist on the same miRNAs. This finding is further supported by the consistent expression patterns observed for these shared miRNAs across both PDAC and NAT tissues. These observations suggest a potential role for these co-glycosylated miRNAs in pancreatic cancer. Finally, downstream target prediction analysis identified potential target genes associated with both the significantly shared and unique miRNAs (**Figure 5D**). These findings provide valuable insights into the functional implications of miRNA glycosylation in pancreatic cancer and warrant further investigation into their specific roles in disease progression.

### Glycosylated miRNAs in PDAC target genes involved in tumor proliferation and cytokine response

We identified 14 miRNAs with significantly enriched target genes in the TCGA-PDAC database. These miRNAs comprise 3 unique and 11 shared miRNAs across PDAC and NAT tissues. **Figure 6A** depicts the high-confidence target genes of these miRNAs: the upper network visualizes those targeted by the shared miRNA targets, while the lower network shows those targeted by the unique miRNA targets. The top enriched Gene Ontology (GO) function terms and the top enriched KEGG pathways are shown in **Figure 6B** and **Figure 6C**, respectively. These target genes revealed distinct functional roles for shared and unique glyco-miRNAs in PDAC. Both sets targeted genes involved in tumor cell proliferation through the PI3K-Akt signaling pathway, which is well known for its role in cell growth and survival ^32^. However, unique glyco-miRNAs focused primarily on processes related to the immune system, including hematopoiesis (blood cell formation), cytokine signaling, and signal transduction pathways. Shared glyco-miRNAs, on the other hand, emphasized target genes may be associated with tumor development, metabolism, and cell adhesion. Overall, these findings suggest that glycosylated miRNAs likely play a significant role in PDAC development and progression. The differential targeting patterns between unique and shared miRNAs hint at the possibility that different glycan modifications on RNA may have distinct effects on its function.

**Figure 6.**
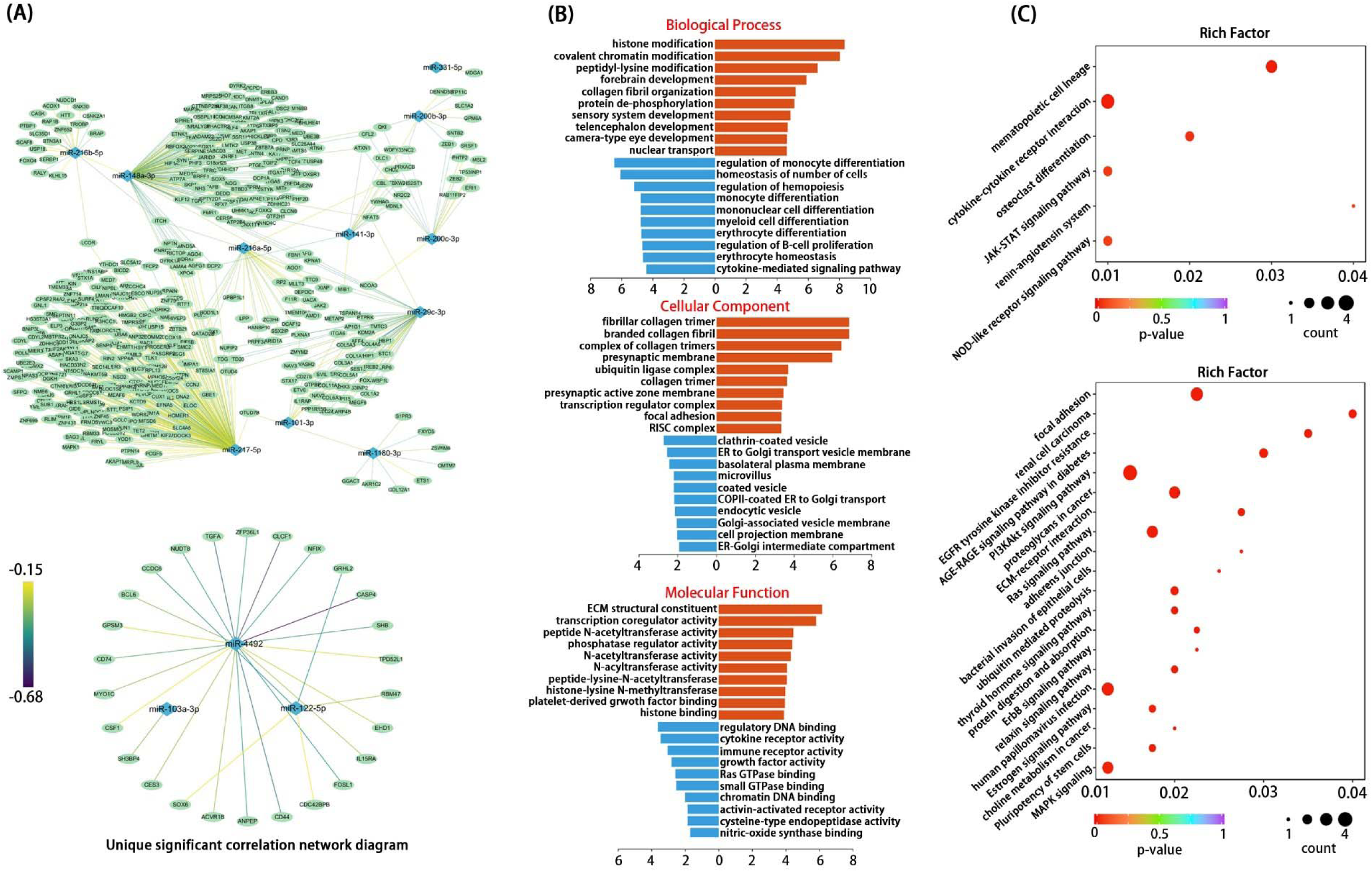
Identification of shared and unique glycosylated miRNA target genes associated with PDAC and their enrichment in biological processes, cellular components, and KEGG pathways. (A) Shared and unique glyco-miRNA targets correlate predicted common target genes (13 glyco-miRNAs and 3 unique glyco-miRNAs) with TCGA-PDAC data (178 cases). The upper network shows significantly correlated genes targeted by shared miRNAs and the lower network by unique miRNAs (1 gene for miR-103a-3p, 5 for miR-122-5p, and 20 for miR-4492, **Table S2**). (B) Gene Ontology (GO) enrichment for biological process (BP), cellular component (CC), and molecular function (MF). Five terms are displayed with - log10(p-value) on the x-axis. (C) KEGG pathway enrichment on 13 significantly shared miRNAs in PDAC. The top 6 significant pathways are shown. KEGG enrichment on 3 unique miRNAs is visualized for the top 15 significant pathways. The PI3K-Akt signaling pathway is enriched for all miRNAs and regulates cell growth, proliferation, and survival. The JAK-STAT signaling pathway is uniquely enriched for the unique miRNAs and is involved in cytokine response.

### PONglyRNA: an online tool for site-specific prediction of glycosylation in RNA

Direct detection of glycosylation sites on RNA molecules remains a significant challenge due to their rapid degradation. However, indirect methods like TnORNA and SPCgRNA offer valuable tools. These methods can confirm the presence of glycosylation on already identified RNAs. Additionally, PONglyRNA provides a complementary approach for localizing potential glycosylation sites within RNA molecules. This localization capability facilitates further investigation into the functional roles of glycoRNAs within cells.

Following data preprocessing and transcript mapping, our results revealed that N-linked and O-linked glycosylation sites were distributed across various RNA types. MiRNAs exhibited the highest proportion of glycosylation sites, followed by snRNAs and snoRNAs (**Figure 7A**). To understand the sequence preferences associated with glycosylated RNAs (glycoRNAs), we identified the most frequent motifs within each RNA category (snRNA/snoRNA, miRNA precursors, tRNA) and compared them to known RNA-binding protein (RBP) motifs. Interestingly, the analysis revealed enrichment of motifs similar to those recognized by specific RBPs in both N-linked and O-linked transcripts (**Figure 7B**&**C**, **S3A**-**D**). For snRNA/snoRNA transcripts, these motifs resembled binding sites for *RBMS3*, *PTBP1*, *ZC3H10*, *PABPN1*, and *ZNF638*; miRNA precursor transcripts displayed enrichment of motifs analogous to those bound by *MSI1*, *A1CF*, *RBM8A*, *PCBP2*, and *SEPQ*; N- and O-linked tRNA transcripts harbored the most enriched motifs corresponding to *RBM4*, *ESRP2*, *EIF4B*, *RBM4B*, *RBM6*, *SNRPB2*, *ZCRB1*, *FUS*, *NONO*, and *SNRP70*.

**Figure 7.**
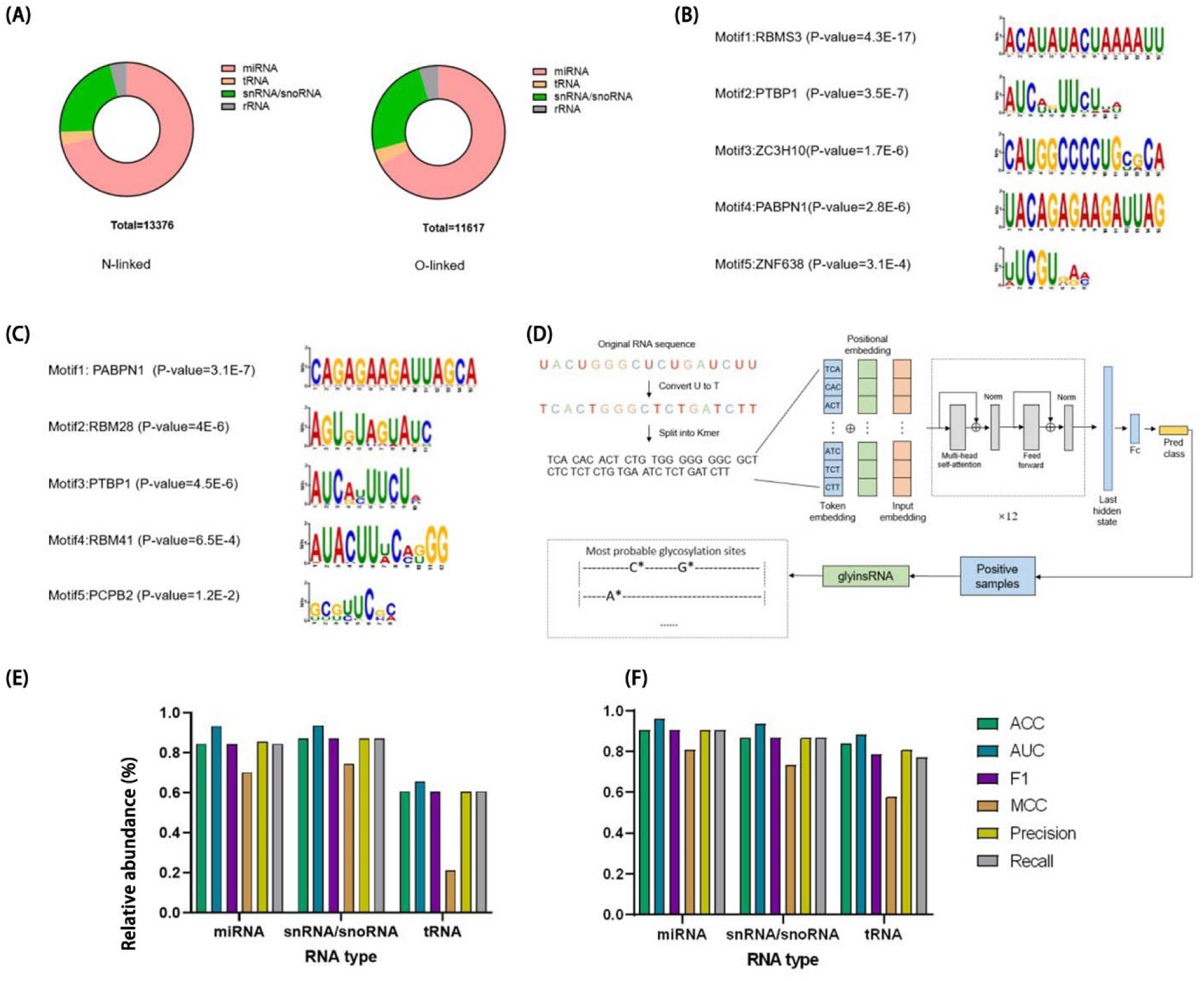
PONglyRNA: A Tool for Predicting O- and N-Linked Glycosylation in RNA. PONglyRNA is a computational tool that predicts the presence and location of O- and N-linked glycosylation sites on RNA molecules. (A) Distribution of RNA types (not RNA abundance) containing predicted O-linked and N-linked glycosylation sites within the PONglyRNA dataset. (B & C) The five most prevalent sequence motifs identified among RNAs predicted to harbor (B) N-linked and (C) O-linked glycosylation, respectively. (D) Schematic representation of the PONglyRNA workflow. (E & F) Performance evaluation of PONglyRNA in predicting the presence of (E) N-linked and (F) O-linked glycosylation on RNA.

The above analysis suggests that RNA sequence features can be used to predict the presence of glycosylation. Therefore, we developed PONglyRNA which utilizes RNA transcript sequences for prediction. **Figure 7D** illustrates the workflow of PONglyRNA. This framework employs a pre-trained DNABERT module, fine-tuned for our specific task, to predict glycosylation sites within full-length sequences of small RNAs (miRNAs, tRNAs, etc.) containing either N-linked or O-linked modifications. By feeding the model full-length sequences with potential glycosylation sites, PONglyRNA learns to extract relevant features from the nucleotide sequence and predict transcripts likely to harbor these modifications. **Figures 7E** and **7F** illustrate PONglyRNA’s performance in predicting RNA N- and O-linked glycosylation, respectively. The model demonstrates strong accuracy in identifying glycosylation potential within miRNA precursors and full-length snRNA/snoRNA sequences. However, the limited number of tRNA samples resulted in lower prediction accuracy for this RNA type. Due to the scarcity of negative rRNA samples, we excluded rRNA from the model development process.

To aid the research community, PONglyRNA is publicly accessible as a user-friendly online web server at http://ponglyrna.797000.xyz:8880/. Users can simply submit their RNA sequence and choose the RNA type for prediction. The platform automatically runs the deep neural network model and provides detailed prediction results, including a visualization of the predicted modification sites. For instance, **Figure S4** showcases the distribution of N-linked glycosylation sites with a probability exceeding 0.6 within a full-length snRNA sample.

## Discussion

The field of glycoRNA research, the latest component of glycome, is rapidly evolving. This field sheds light on a new layer of complexity in cellular regulation: RNA glycosylation, the process of attaching glycans to RNAs. This process empowered novel glycan-dependent regulatory role of RNAs. Similar to how glycans act as recognition tags on proteins, they can potentially influence RNA stability, folding, localization, and interaction with other molecules, opening doors to unexplored regulatory pathways within the cell. Unlike protein glycosylation which is limited to specific locations, RNA can potentially be decorated with glycans throughout the molecule. This broader modification suggests that RNA glycosylation might be involved in a wider range of cellular processes beyond just protein production. Compared to protein glycosylation, RNA glycosylation presents exciting new possibilities. Glycans might act as cellular GPS (global positioning system), directing RNA molecules to specific compartments and influencing their function. This is particularly relevant for ncRNAs that lack protein-coding abilities, where glycosylation could be a unique mechanism to control their activity ^33^. Understanding how glycosylation affects RNA function could pave the way for novel therapeutic strategies targeting specific RNA molecules in diseases. However, the field remains in its infancy. RNA glycosylation lags far behind the extensive knowledge base on protein/lipid glycosylation. The specific functions and mechanisms by which RNA glycosylation operates are yet to be fully elucidated.

This work establishes a robust solid-phase chemoenzymatic approach for specifically targeting O-linked RNA glycosylation. The key to capturing O-glycoRNAs lies in the presence of Galactose (Gal) and/or N-Acetylgalactosamine (GalNAc). These monosaccharides can be oxidized by the enzyme GAO. TnORNA, which utilizes GAO, can theoretically oxidize most O-glycans except for core 3 and core 4 structures (**Figure S2**). Other types of O-glycans can be oxidized by either Gal or GalNAc. Importantly, GAO cannot oxidize Gal or GalNAc capped with sialic acids linked at the C-2 or C-6 positions (α2-3 or α2-6 linkages) because the C-6 position is occupied by the sialic acid ^34^. However, sialic acid residues on Gal or GalNAc can be first removed by the enzyme sialidase (SiaEXO). This exposes the C-6 position, allowing subsequent oxidation by GAO ^34,35^. Therefore, TnORNA, in combination with sequential sialidase treatment, can tag O-glycoRNAs containing sialyl T or sialyl Tn structures as well.

TnORNA successfully enriches various ncRNAs, including miRNA, tRNA, rRNA, snRNA, and snoRNA, suggesting a broad role for O-glycosylation in diseases (**Figure 3**). While current research prioritizes glycosylated miRNA, targeting other O-glycoRNAs like modified rRNAs remains a promising avenue for future treatment strategies. rRNAs undergo extensive modifications during biogenesis, with these changes strategically placed to stabilize the ribosome structure and facilitate protein synthesis ^36,37^. These modifications are installed by either snoRNA-guided or standalone enzymes within the nucleolus, nucleus, and cytoplasm. Recent discoveries of sub-stoichiometric 2’-O-methylation and pseudo-uridylation challenged the previous notion of complete rRNA modification ^38,39^. These findings highlight the role of diverse alterations in generating ribosome heterogeneity. The mechanisms regulating partial modification and the functional specialization of ribosomes remain largely unknown. However, observed changes in rRNA modification patterns in response to environmental cues, development, and disease suggest a potential role in dynamically controlling gene expression through translation. Further investigation into snoRNA glycosylation is crucial to understand its potential impact on both rRNA structure and function.

TnORNA and SPCgRNA offer complementary methods for enriching distinct types of glycoRNAs: TnORNA targets O-linked glycosylation, while SPCgRNA enriches for N-glycoRNAs. Applying these techniques to MIA PaCa-2 cells and pancreatic tissue samples revealed the presence of both N-linked and O-linked glycoRNAs within the cells (**Figures 4** & **5**). Interestingly, some RNA molecules even displayed co-occurrence of both N- and O-linked modifications. Our data suggests a positive correlation between the expression of specific miRNAs and their glycosylation level. This dysregulation of glycosylation appears particularly significant in pancreatic cancer for miR-200b-3p, miR-148a-3p, miR-1180-3p, and miR-216b-5p. Notably, miR-200b-3p functions as a well-established tumor suppressor by inhibiting metastasis. Studies have shown decreased levels of this miRNA in hepatocellular carcinoma tissues, while its overexpression can reduce cell migration and proliferation ^40^. Similarly, in glioblastoma patients, adenosine methylation of miR-200b-3p disrupts its ability to suppress target mRNAs like XIAP (X-linked inhibitor of apoptosis protein), potentially contributing to poorer clinical outcomes ^41^. These findings collectively suggest that miRNA glycosylation likely modulates their functions in diseases. However, further investigation is necessary to pinpoint the exact locations and specific nucleotides undergoing glycosylation on these miRNA molecules.

While TnORNA and SPCgRNA respectively enrich for O- and N-glycoRNAs, they don’t pinpoint the exact glycosylation sites on the RNA molecule. This limitation arises from the highly fragile nature of RNA, making site-specific analysis extremely challenging with current techniques. However, since our methods can identify which RNAs are glycosylated, we can leverage these sequences for site-specific prediction. Here, we employ RNA sequences to predict potential glycosylation sites and their associated motifs. This allows us to identify the likelihood of glycosylation at specific locations. By site-specific mutations based on these predictions, we can investigate the role of RNA glycosylation in disease development and progression.

This study explored the target genes of glycosylated microRNAs (miRNAs) in pancreatic ductal adenocarcinoma (PDAC). Analyzing TCGA-PDAC data, we identified shared and unique sets of miRNAs with significantly correlated target genes (**Figure 6**). These target genes were enriched in biological processes crucial for cancer progression, including cell proliferation and cytokine response ^42^. Furthermore, KEGG pathway analysis revealed significant associations with established cancer pathways, such as the PI3K-Akt pathway (regulating cell growth and survival) and the JAK-STAT pathway (involved in growth factor response). These findings suggest that glycosylated miRNAs target genes involved in key hallmarks of cancer, potentially influencing PDAC development and progression. Notably, both shared and unique miRNA sets impacted pathways critical for tumorigenesis, highlighting their potentially diverse functionalities. The identified associations with the PI3K-Akt and JAK-STAT pathways warrant further investigation in pre-clinical models to elucidate the mechanisms by which glycosylation modulates miRNA function and target selection. Ultimately, this study paves the way for exploring the therapeutic potential of targeting glycosylated miRNAs in PDAC. **Acknowledgments**

This work was supported by the Soochow University Start-up Fund, the Priority Academic Program Development of the Jiangsu Higher Education Institutes (PAPD), Jiangsu Science and Technology Plan Funding (BX2022023), the Jiangsu Shuangchuang Boshi Funding (JSSCBS20210697), Suzhou Project Funding (SKY2021026), Suzhou Medical Innovation Funding (SKJY2021141), Pilot Research Funding (SDFEYGJ2001) of the Second Affiliated Hospital of Soochow University and Open Project (GZK12023012) of State Key Laboratory of Radiation Medicine and Protection, National Natural Science Foundation of China (32070658 and 32222020).

## Author Contributions

J.J.L.: Performed all experiments, drafted the experimental methods section, and analyzed the data. S.Y.: Designed the experiments, prepared the figures and tables, and drafted the manuscript. L.W. and Y.Z.: Developed PONglyRNA and drafted the manuscript. Y.C., S.Z., and L.J.X.: Collected clinical specimens and participated in sample preparation. Z.M.W., H.Y.Z., L.J.X., Y.Z. and S.Y.: Secured funding for the research. All authors: Approved the final manuscript.

## Methods

### Cell culture of PDAC cell lines

The human pancreatic ductal cancer cells, MIA PaCa-2, and human non-small cell lung cancer cells, A549, used in this experiment were purchased from Wuhan Procell Life Technology Co., Ltd. Their short tandem repeat identification was confirmed. MIA PaCa-2 cells were cultured in DMEM medium containing 1% double antibody (penicillin-streptomycin) and 10% fetal calf serum. A549 cells were cultured in RPMI-1640 medium containing 1% double antibody and 10% serum. All cells were cultured in a cell culture incubator at 37°C and 5% CO_2_.

### Total RNA extraction from cells

Once the cells reach confluency in T75, T25, or six-well plates, they were harvested for further processing. The culture medium was replaced with fresh, ice-cold 1xPBS (repeated twice). TRIzol reagent (1 mL per T75 flask) was then added to the harvested cells and lysed at room temperature for 5 minutes. A cell scraper was used to collect the lysed cells, which were then transferred to an enzyme-free, sterile centrifuge tube. The cell lysate was incubated on ice for 5 minutes to allow for phase separation. Following stratification, the upper aqueous phase, containing RNA (pH < 7), was carefully transferred to a new enzyme-free centrifuge tube. The remaining pellet was washed with 75% ethanol (tube briefly inverted) and centrifuged at 8,000 rpm for 5 minutes at 4°C. This washing step was repeated three times. The washed pellet was dried upside down for approximately 5 minutes. Finally, 20 µL of DEPC water was added to dissolve the pellet, resulting in the total RNA solution.

### Total RNA extraction from FFPE

#### Removal of FFPE wax by mineral oil

The processing steps depend on the tissue slice thickness. For slices less than 50 μm, 300 μL of mineral oil was added; for thicker slices (>50 μm), 500 μL of mineral oil was used. Following oil addition, the sample was incubated in a metal bath at 80°C for 1 minute and then vortexed to mix thoroughly. 100 μL of lysis buffer was added, and the sample was centrifuged at room temperature (10,000 × g for 15 seconds). This procedure can separate the sample into an upper oil layer and a lower water layer. Proteinase K (10 μL) was directly added to the lower (blue) water layer and mixed thoroughly with a pipette. The sample was then incubated in a 56°C metal bath for 15 minutes to ensure complete tissue dissolution. During incubation, the two layers should be mixed thoroughly by inversion. If complete dissolution was not achieved, we extended the incubation time to 30 minutes. Finally, the sample underwent further incubation in a metal bath at 80°C for 1 hour. After removal from the 80°C bath, the sample was cooled on ice for 1 minute and then brought to room temperature for 2 minutes.

#### DNase treatment

30 μL of fresh DNase treatment mixture was added to the water layer. The water phase was then gently mixed using a pipette. Following this, the sample was incubated at room temperature for 15 minutes.

#### Nucleic acid binding

A mixture of 325 μL LB buffer (lithium borate) and 200 μL of 100% isopropanol was gently mixed. This mixture was then centrifuged at room temperature (10,000 x g for 15 seconds). Following centrifugation, the upper oil layer was carefully separated from the lower water layer. For each sample, an adsorption column was placed in a corresponding collection tube. The entire blue aqueous phase was carefully transferred to the adsorption column with its cap still closed. The remaining oil layer was discarded. Finally, the collection tube containing the adsorption column was centrifuged again at room temperature (10,000 x g for 30 seconds). The waste from this centrifugation was stored, and the clean adsorption column was transferred to a new collection tube.

#### Washing and elution

500 μL of 1x wash buffer was added to the adsorption column. (Note: Ensure ethanol was added to the wash buffer before use.) The column cap was then closed, and the sample was centrifuged at room temperature (10,000 x g for 30 seconds). The discarded waste liquid was followed by transferring the adsorption column to a new collection tube. This washing step was repeated once. Subsequently, the capped adsorption column was further dried by centrifugation at 16,000 x g for 3 minutes. The column was then transferred to a new 1.5 mL centrifuge tube and allowed to sit at room temperature for 5 minutes. Next, 50 μL of RNase-free water was added to the column, and the cap was closed for incubation at room temperature for 5-10 minutes. Finally, the RNA solution was obtained by centrifugation at room temperature (16,000 x g for 1 minute).

### TnORNA method

Building upon previously described total RNA extraction, this procedure detailed the processing of the resulting total RNA solution (fraction #1) to obtain oxidized O-glycosylated RNA. Galactose oxidase (GAO) was employed to oxidize the hydroxyl group of either Galactose (Gal) or N-Acetylgalactosamine (GalNAc) residues into aldehyde groups. The specific steps were as follows: 40-60 μg of total RNA is mixed with 20 μg each of galactose oxidase (GAO) and horseradish peroxidase (HRP) in 20% DMSO. This reaction mixture was then incubated for 2 hours at room temperature in the dark.

#### Solid-phase glycoRNA immobilization

Following oxidation, the RNAs were covalently immobilized onto beads containing hydrazide groups. This process involved mixing 100 μL of hydrazide bead slurry with the oxidized RNA solution and incubating the mixture at room temperature for 2 hours. This allows the glycan-containing RNAs to bind to the hydrazide beads, while non-glycosylated RNAs remain unbound. Unbound RNA was subsequently removed through a washing step. The sample was rinsed with diethyl pyrocarbonate (DEPC) water, and this washing process was repeated three times. The final washing solution (fraction #2) was collected, and its RNA concentration should be measured. Ideally, this measurement should be zero, indicating the successful removal of non-glycosylated RNA. Following confirmation of this step, the protocol could proceed to the next stage.

#### O-glycoRNA enzymatic release

Following immobilization, the O-glycans on the RNA were specifically digested using either O-glycosidase (targeting core 1 and core 3 structures) or N-acetylgalactosamine hydrolase (GalNAcEXO) for Tn-antigen containing O-glycans. For Tn-antigen digestion, 100 μL of 20 mM ammonium bicarbonate (NH_4_HCO_3_) buffer was first added to the RNA-conjugated beads, followed by 20 μL of GalNAcEXO. This enzymatic reaction proceeded at room temperature for 90 minutes, resulting in the elution of O-glycosylated RNA (fraction #3). Similarly, 20 μL of O-glycosidase was added to a separate aliquot of the beads and incubated for 90 minutes to target different core structures (fraction #4). After enzymatic treatment, the eluted fractions (#3 or #4) were then subjected to further analysis, including concentration measurement, absorbance measurement, gel electrophoresis, and high-throughput RNA sequencing.

#### TnORNA enrichment specificity

To evaluate the specific effects of enzymes and MUC5AC on Tn-containing RNA (TnORNA), we performed experiments with GalNAcEXO and SiaEXO + GalNAcEXO, both in the presence and absence of MUC5AC, prior to the GAO oxidation step. This approach aims to assess the influence of GalNAc, sialic acid (Sia), and MUC5AC on TnORNA. For GalNAc removal, 20 mM NH_4_HCO_3_ buffer was added to the sample, followed by incubation at 25°C for 2 hours. To remove sialic acid, 40 μL of SiaEXO was added to the 20 mM NH_4_HCO_3_ buffer, and the mixture was incubated at 25°C for 2 hours. Finally, the effect of the O-glycoprotein was tested by adding 1 μg and 12 μg of MUC5AC to the sample.

### PONglyRNA for site-specific prediction of glycoRNA

#### PONglyRNA data processing and prediction algorithm

Raw N- and O-linked glycoRNA reads were quality controlled using the fastp tool ^43^. After filtering, the sequences were aligned to reference RNA databases (RefSeq for mRNAs and ncRNAs, Rfam for ncRNA families, and miRbase for pre-miRNA hairpins) using the blastn-short tool with stringent parameters: 100% identity (perc_identity), word size of 7, and e-value threshold of 1. Full-length RNA sequences with at least one mapped glycoRNA read were designated as positive samples. An equal number of transcripts of the same RNA type from the reference databases were selected as negative examples. To remove redundancy within the dataset, CD-HIT-EST was employed with a 0.9 sequence identity threshold ^44^. Next, the full-length RNA sequences were encoded using DNABERT, a pre-trained Transformer model capable of capturing both global and local sequence information ^45^. The encoded sequences were fed through a transformer layer followed by two fully connected layers to train the glycoRNA classification model. Hyperparameters were optimized during training, including a learning rate of 3 × 10^-5^ and a warmup percentage of 0.1. Finally, the predicted glycoRNAs were analyzed by our previously developed GlyinsRNA algorithm to pinpoint the glycosylation sites ^46^.

#### Motif analysis

To identify potential RNA-binding protein (RBP) interactions, we utilized a motif discovery and comparison approach. We employed the MEME Suite’s STREME and Tomtom tools ^47,48^. First, STREME identified statistically over-represented sequence motifs in the positive samples compared to the negative samples. The search focused on motifs with widths ranging from 8 to 15 base pairs (bp) and applied a p-value cutoff of 0.05 to ensure statistical significance. Subsequently, Tomtom was used to compare these over-represented motifs against known RBP motifs deposited in the RNAcomplete and CISBP-RNA databases ^49^. This comparison allows us to identify potential RBPs that might interact with the RNA sequences under study.

## Supplemental Information

Additional experiments, including comparison of digestion efficiency, identification of small RNA sequence, small RNA data processing, and RNA data processing, are listed in the supporting materials. Figure S1 – TnORNA optimization; Figure S2 – GAO oxidation; Figure S3 – glycoRNA motif; Figure S4 – PONglyRNA prediction; Table S1 – glycosylated miRNA; Table S2 – glycosylated miRNA correlation.

